# Simple and Multiplexed Tracking of Bacterial Growth in Double Emulsion Droplets

**DOI:** 10.64898/2026.05.01.722333

**Authors:** Sebastián Somolinos Cedeño, Samuel Thompson, Polly Fordyce, Drew Endy

## Abstract

Droplet-based microfluidics enable researchers to observe phenotypic heterogeneity within complex biological mixtures through parallel encapsulation of individual samples followed by imaging. Observing or quantifying dynamic heterogeneity remains challenging due to complexities associated with trapping and tracking many individual droplets. Current approaches for time-lapse imaging require specialized devices with droplet traps that limit accessibility and throughput. Here, using readily available materials and software, we demonstrate a simple method for stabilizing and monitoring many, individual droplets for up to 12 hours. We leveraged our method to track bacterial growth within droplets in a high-throughput manner. Our method allows tracking the changes and variation in growth rate within and across droplets, revealing heterogeneity in growth patterns hidden in batch assays. Improving the affordability and throughput of time-dependent phenotyping assays helps to advance biological discovery and biotechnology innovation.

## Introduction

Droplet microfluidic technologies enable high-throughput imaging of individuals comprising complex populations (1–5). Via highly parallelized compartmentalization, individuals suspended in a continuous aqueous phase can be isolated and encapsulated within an external oil shell with high precision and speed (6), creating many distinct and independent samples.

Double-Emulsion (DE) droplets have gained attention relative to single-emulsion (SE) approaches (7–9). DE droplets are composed of an inner aqueous phase, an oil shell, and an outer aqueous phase (Fig. 1A). Surfactants stabilize each water-to-oil interface, lowering the surface tension to allow the droplets to self-assemble in a microfluidic generator and providing sufficient repulsion to protect droplets from coalescing. The outer aqueous phase further reduces the likelihood of droplet merging and allows for nutrient exchange. DEs also enable droplet isolation through Fluorescence Activated Cell Sorting (FACS), circumventing the complex set-ups required for sorting single-emulsion droplets through Fluorescently Activated Droplet Sorting (FADS) (10).

**Figure 1:**
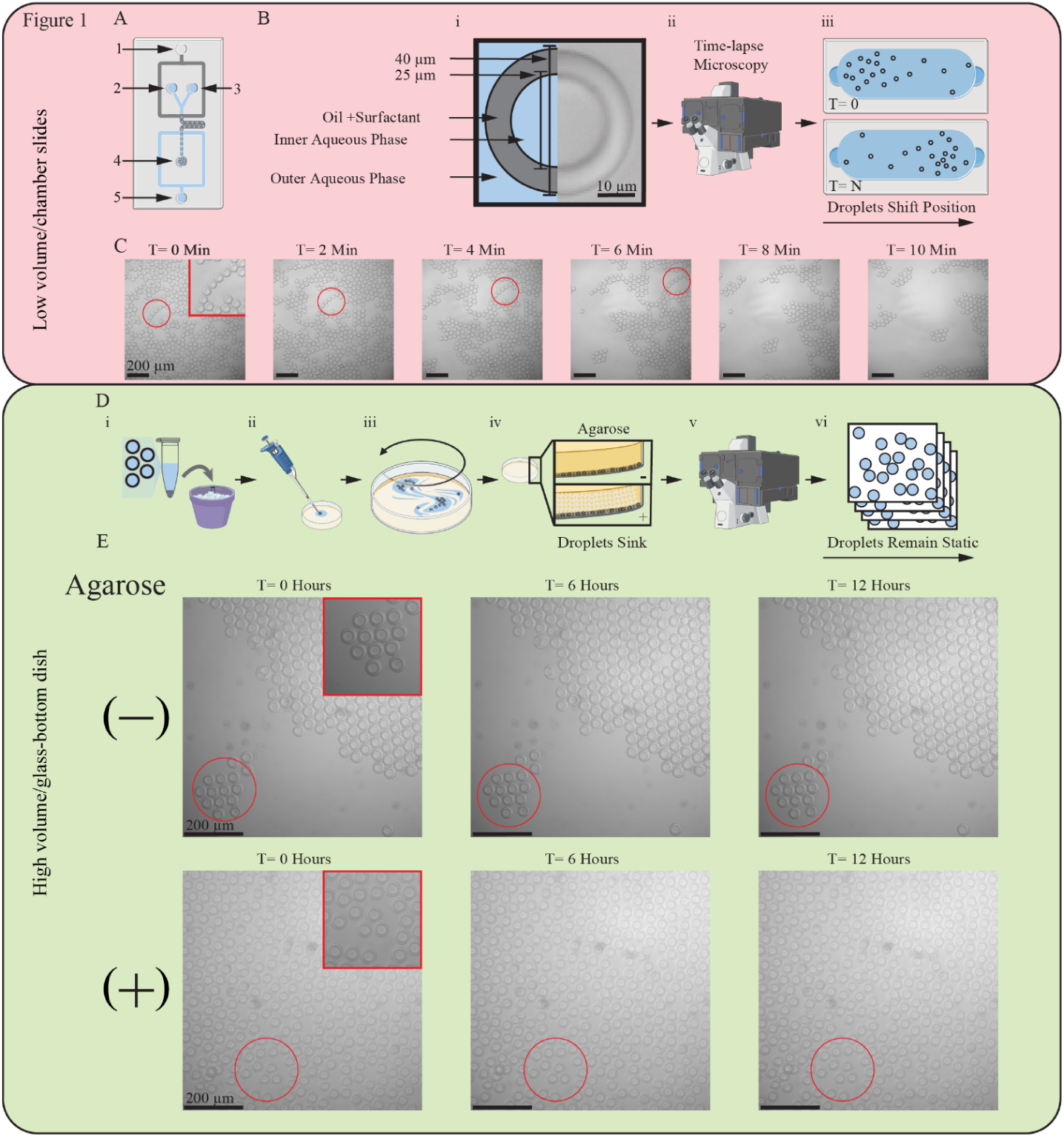
Agarose trapping enables long-term tracking of double emulsion (DE) droplets. (A) Schematic of DE-droplet generator device. (B) Schematic showing the problem addressed in this work: 1) DE-droplets (layers and dimensions included) are generated, 2) introduced into a temperature-controlled microscope, and 3) lost during imaging due to positional shifting within the chamber slide. (C) Bright field microscopy images of a single field of view (FOV) containing DE-droplets over time; Red circle indicates the position of a particular set of droplets as they migrate. (D) Schematic of experimental pipeline for immobilizing DE droplets in agarose followed by time-lapse imaging: 1) generate droplets and keep on ice until imaging, 2) inject droplets at the bottom of a glass microscopy dish filled with molten agarose, 3) gently swirl the dish, 4) allow agarose to solidify for 5 minutes, 5) place dish on the microscope stage, and 6) perform time-lapse imaging of desired FOVs. (E) Bright field microscopy images showing droplets immobilized using different agarose concentrations at 0 hours (left) and 12 hours (middle) and particle tracking data of per-droplet movement (right).

Researchers have realized a wide variety of enzymatic and cellular screens by sorting microfluidic droplets to isolate and recover individual samples expressing desired properties (11–18). DE droplets specifically have been used to isolate and enrich diverse biological entities including DNA molecules, enzymes, viruses, and prokaryotic and eukaryotic cells (19–23). Partitioning populations into individual compartments enables growth in the absence of competition, facilitating measurement of low-fitness or uncommon variants that would otherwise be masked by faster-growing subpopulations if tested in bulk (24).

DE-droplets can also be useful during imaging. Microscopy of droplets can provide quantitative information about time-dependent processes taking place within thousands of individual droplets in parallel (25). Heterogeneous behaviors that are obscured when averaging across droplets or limiting data collection to static imaging can also be revealed (26). Parallel microencapsulation of individuals enables access to the diversity of phenotypes underlying population-level measurements, revealing insights into the dynamic range of certain kinetic processes. Continuous monitoring of specific droplets could also help screen enzyme variants for turnover speed, or help determine the prevalence of a sub-population of growth-deficient cells. Beyond real-time observations of emergent dynamic heterogeneity, time-lapse DE droplet imaging could provide information critical in the process of developing, troubleshooting, and improving novel high-throughput screens (27,28). Taken together, simple and scalable pipelines for tracking DE droplets could expand the accessibility and usefulness of high-throughput imaging for complex biological samples (29).

Tracking droplet identity and phenotype over time remains challenging because droplets tend to move, shifting in and out of the field of view and focal plane (29). Because droplets move in response to external forces (e.g., evaporation, jostling) existing approaches often rely on repeated loading of sample aliquots at different time points. While static imaging can capture population-level measurements, doing so precludes longitudinal tracking of individual droplets, which is essential in understanding the underlying causes of population dynamics and heterogeneity (26). Collecting droplet tracking data should ideally be simple, with the throughput only limited by the size and number of captured fields of view (FOV).

Previous studies performing time-lapse imaging of droplets have required specialized secondary devices containing droplet traps (2,5,30–35). Designing and operating devices with droplet traps is challenging, often requiring fabrication of new multi-height devices with tailored trap geometries (26) and either continuous flow (36) or valves (37) for operation. The manufacturing process for such devices requires expertise, access to a clean room, and often several rounds of iteration during fabrication. Additionally, most computational droplet tracking methods rely on custom software, complicating accessibility and reproducibility. Custom tracking algorithms are often tailored to a particular use case and can be computationally expensive, taking up to four days to process an experimental run (31). Other methods using broadly-accessible software rely on secondary trapping devices and have yet to be applied to tracking of DE droplets (2).

We developed a simplified imaging and tracking pipeline to streamline multiplexed monitoring of DE droplets. We only used readily-available consumables and software to help reduce barriers to future adoption. As a benchmark, we monitored the increase in bacterial cell density within droplets over time, expecting an initial lag to be followed by an exponential increase, and ultimately becoming stationary. Plotting the resulting data as a line chart revealed a sigmoidal curve typical of bulk-based assays. Beyond recapitulating expected cell behaviors, our method enabled tracking of bacterial growth within hundreds of DE droplets in a single FOV over several hours, and processing of the resulting data within minutes. Furthermore, inspired by the agarose pad method developed by Elowitz to observe monolayers of growing and dividing bacteria (38), we demonstrated compatibility of DE droplets with agarose trapping during high-throughput imaging and tracking.

## Results

### Use of a glass-bottom dish, increased sample volumes, and agarose help minimize droplet drift

To demonstrate the severity of the droplet shifting issue in trap-less imaging set-ups, we tested how long we could track and image DE droplets without immobilization. Droplets generated using a standard PDMS microfluidic chip (Fig. 1A) were roughly 40 µm in diameter and contained a ∼25 µm inner core of PBS surrounded by a shell of HFE7500 oil with 5% PicoSurf surfactant (Fig. 1B, i). We generated a monolayer of DE droplets suspended in PBS by depositing them on a Countess chamber slide (Thermo Fisher Scientific), and then imaged droplets via time-lapse, bright-field microscopy at two-minute intervals (Fig. 1B, ii). Images of droplets showed substantial displacement over time, with droplets from a packed field of view gradually moving out of the FOV (Fig. 1B, iii).

The speed and magnitude of displacement was severe enough to preclude computationally tracking the identity of any given droplet, even at high temporal resolution. Most droplets present in the first frame (t = 0) drifted out of the original FOV by the fifth frame (t = 10 minutes) (Fig. 1C).

We pursued two solutions to limit droplet movement. First, we attempted an optimized “trap-less” method that kept droplets unconfined. To prototype a simple implementation of a trap-less imaging set-up, we used a glass bottom dish (D35C4-20-0-N, Cellvis) and increased the relative volume of the outer buffer (Fig. 1D). We reasoned that using glass would help reduce shifting due to temperature-induced surface bending and increased buffer volume would reduce the impact of evaporation. The high density of DE-droplets ensured they would sink and form a monolayer at the glass interface, reducing vertical movement while maximizing imaging throughput. Second, we tested the compatibility of DE droplets with agarose-based trapping (Fig. 1D, iv and Supp. Fig. 1). We hypothesized that droplets embedded in agarose would remain fixed in a FOV for the duration of a growth experiment without requiring additional device fabrication. We added agarose at 0.25 to 1% w/v into our outer buffer (PBS). Through microscopy, we observed consistent stability in droplet integrity, focal plane, and relative position within the FOV for up to 12 hours, regardless of agarose presence (Fig. 1E and Supp. Fig. 2).

Our results suggest that DE droplets are compatible with agarose trapping and that either pipeline was robust enough to prevent excessive droplet shifting for up to 12 hours. Taken together, our findings suggested downstream compatibility with multiplexed imaging and tracking of time-dependent processes in DE droplets, such as bacterial growth.

### Fluorescence Imaging of DE-droplets Reveals Differences in Cell Density

Next, we were curious about the sensitivity of our simplified, single-plane imaging set-up and whether it could detect differences in cell densities well enough to detect growth. At low cell densities, individual bacteria could be difficult to detect inside droplets; at high cell densities, it could be challenging to quantify changes in the absolute number of bacteria. As a proof of concept, we experimentally mocked-up expected conditions at the start and end of a bacterial growth experiment by encapsulating cultures at medium (OD = 0.84) and high (OD = 9.4) density and acquiring a single static image (Fig. 2 and Supp. Fig 3). We selected these cell densities because an OD of 0.84 would lead to a distribution of cells ranging from 0 to 4 cells per droplet (CPD), a range chosen to be easily countable and amenable to the detection of single cells. We choose an OD of 9.4 for a high level of cells (10 to 40 per droplet) because this density would result from three to four cell doublings yet remain well below the theoretical maximum packing of cells per droplet (less than 8000 cells per droplet).

**Figure 2:**
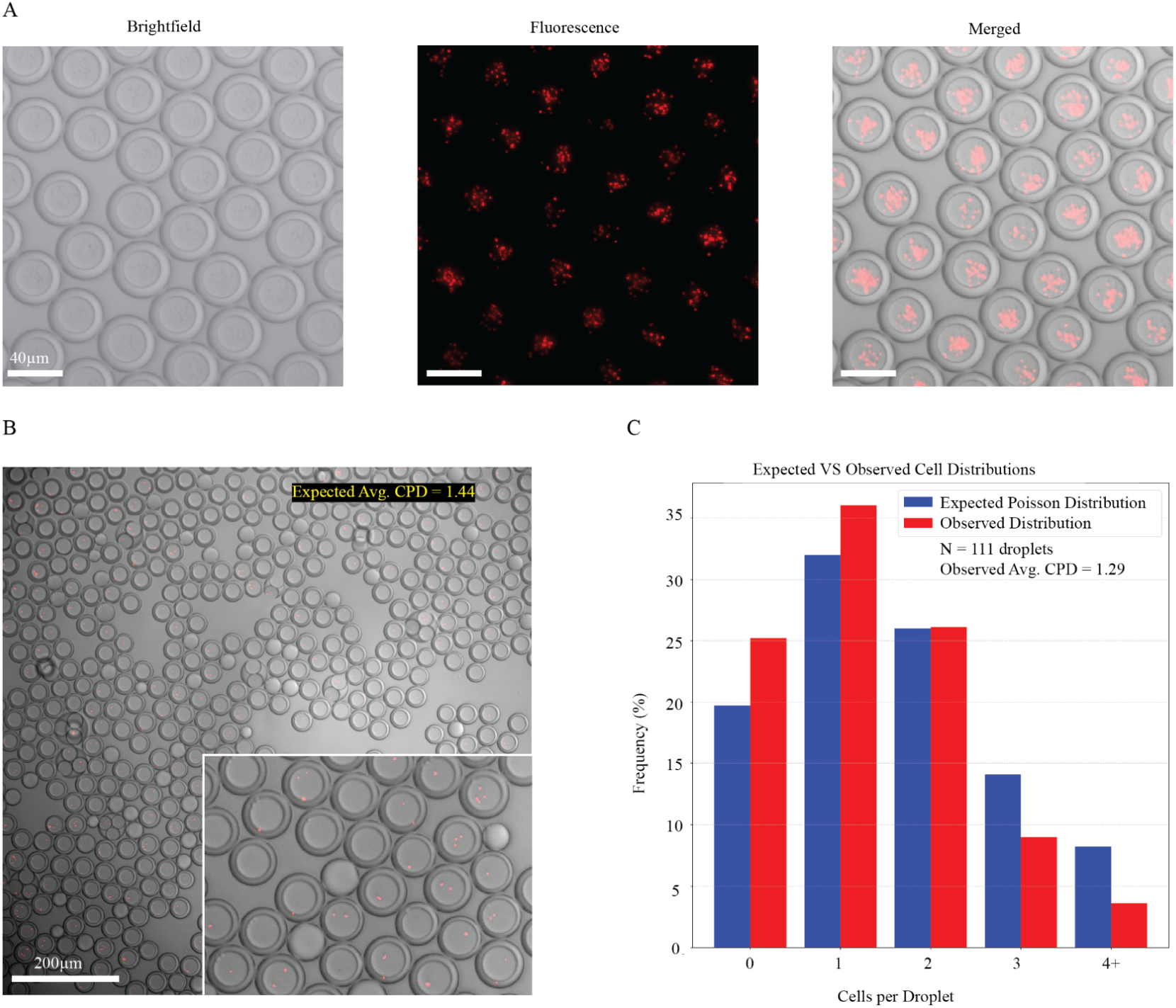
Fluorescence imaging of DE-droplets can reveal differences in the number of cells per droplet. (A) Representative bright field (left), fluorescence (middle), and merged (right) images of a single DE droplet containing mScarlet1-expressing *E. coli* cells loaded under the medium density regime. (B) Example merged image showing a single field of view and zoomed in view (inset) of DE droplets containing mScarlet1-expressing *E. coli* cells loaded under the medium density regime. (C) Histograms showing expected (blue) and observed (red) distributions of the number of cells per droplet.

We encapsulated bacteria in DE droplets by introducing LB-grown cultures and LB alone into the first and second aqueous inner inlets of the microfluidic chip, respectively. The encapsulated *E. coli* constitutively expressed a fluorescent marker (mScarlet1), allowing us to observe cell presence and relative abundance by quantifying pixel intensities (Fig. 2A). Discerning single cells in the high-density regime proved challenging, yet the presence of distinct puncti within medium-density droplets suggested we could resolve individual cells (Fig. 2B).

To further explore the possibility of single-cell detection, we manually counted cells within 111 medium-density droplets and compared our findings to the expected distribution of CPD assuming stochastic loading (Fig. 2C). The averages (λ) of both the observed and expected Poisson distributions within medium density droplets agreed with one another (expected λ = 1.44, observed λ = 1.29 ± 0.1 (SE)). The observed distribution was consistent with the expected Poisson model under a likelihood-ratio test (X^2^(1) = 1.62, p = 0.20).

Our findings showed that our pipeline enabled precise encapsulation and tuning of bacterial density within droplets. The agreement between the observed puncti distribution and the expected cell distribution within medium-density droplets suggested we could indeed resolve single cells. Taken together, our results also suggested that by starting at low cell densities, we could detect and quantify growth of fluorescently labeled *E. coli* during time-lapse imaging, making multiplexed tracking of growth within individual droplets possible.

### Simplified Multiplexed Growth Tracking Reveals Sigmoidal Growth Profiles and Heterogeneity in Growth Rates

We next attempted to track growth of *E. coli* within individual droplets embedded in agarose. We generated droplets using an LB-grown culture at OD of 1, expecting an average of 3.65 cells per droplet. The proportion of empty droplets at T= 400 minutes to total detected droplets at T = 0 (11/370, 2.9%) approximated the expected proportion of empty droplets based on Poisson statistics (2.6 %).

We tracked all detected droplets and recorded data only from those in which the tracking software successfully recorded pixel intensity at all timepoints (N = 299). Pixel intensities within droplets increased over a period of six hours, consistent with cell growth (Fig. 3A, B). Using FIJI’s TrackMate software, we recorded the mean pixel intensity value within every detected droplet within a single FOV over six hours. Mean pixel intensities for most droplets followed a common sigmoidal shape, consistent with expected growth profiles (Fig. 3C). Averaging mean pixel intensity values for all droplets at 5-minute timepoints also generated a sigmoidal curve resembling OD over time plots for batch bacterial cultures moving through lag, log, and stationary phases. On average, the exponential growth phase occurred roughly between 125 and 200 minutes into the experiment, with the maximum average growth rate across all droplets (0.82 doublings/hour) occurring at T = 155 minutes (Fig. 3D)

**Figure 3:**
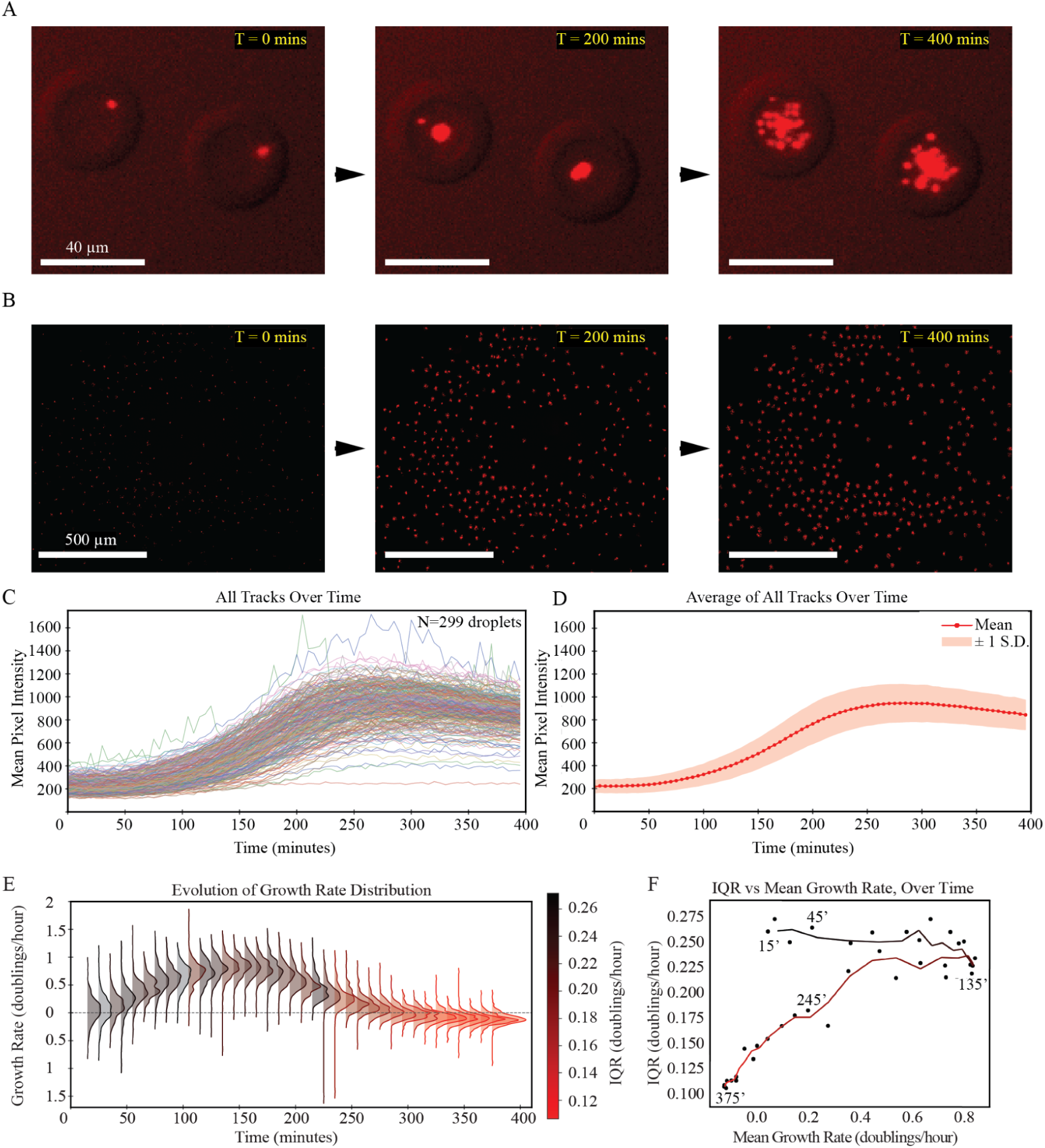
Quantitation of growth rate among single droplets reveals steady and then decreasing variation in growth rate as a function of culture phase. (A) Representative zoomed-in fluorescence images of mScarlet1-expressing *E. coli* within 2 droplets at 0, 200, and 400 minutes. Increase in pixel intensity over time is consistent with cells growing and dividing inside droplets. (B) Representative fluorescence images of mScarlet1-expressing *E. coli* within all droplets within a single FOV at 0, 200, and 400 minutes. Agarose trapping enables detection and tracking of cell growth in many droplets in parallel. (C) Mean pixel intensities for 300 individual droplets over time; each droplet is shown in a different color. Tracking changes in pixel intensity reveals growth profiles consistent with expected sigmoidal growth. (D) Average mean pixel intensities for 300 droplets over time; red line indicates mean and shaded area indicates ± 1 standard deviation of all growth tracks resembles the expected sigmoidal curve from batch assays. (E) Growth-rate PDF time-series colored by IQR (spread of middle 50% of the data), showing a progressive narrowing of the spread in the growth-rate distributions. Dashed line represents a growth rate of 0 (stationary phase) (F) Scatterplot of IQR as a function of mean growth rate over time; Each point represents a single timepoint (15–375 min, 10-min intervals; *n* = 37) from a FOV containing 299 tracked droplets, plotted as IQR (y-axis, doublings/hour) versus mean growth rate (x-axis, doublings/hour). IQRs were computed across all droplets within the FOV at a given timepoint. Lines connect timepoints using a 3-point smoothing to illustrate temporal trajectories, with color transitioning from black (early) to red (late). Time annotations denote representative timepoints (minutes). Across the time course, IQR remains mostly constant until maximal average growth rate is reached, and then decreases as growth begins to slow down.

Next, we wondered if our method could reveal growth rate heterogeneity. Because each droplet serves as an independent, parallel culture, differences in growth rate across droplets might reveal such heterogeneity. We calculated the growth rates (doublings per hour) within individual droplets at a given time point by averaging across 30 minute windows. We then plotted growth rate probability density functions (PDF) in 10 minute increments. We observed an initial increase in the mean growth-rate over time followed by a convergence towards zero in the stationary phase (Fig. 3E). We also noticed a decrease in both mean pixel intensity and growth rate, with the latter becoming negative at late timepoints.

Variation in growth rate across droplets held steady as cells started growing, then decreased as cells stopped growing. Comparing the interquartile range (IQR) between “early growth” (0-115 minutes) and “late growth” (165-275 minutes) periods with equal growth rates (0.41 doublings/hour) revealed a steady decrease in the variance of growth rates over time (early average IQR = 0.25 vs late average IQR = 0.20) (Fig. 3E). Such differences were amplified when comparing individual timepoints with similar growth rates. We observed a 44% increase in IQR across early and late points with roughly equal growth rates of ∼0.20 doublings per hour. (IQR of 0.26 at 45 minutes vs IQR of 0.18 at 245 minutes) (Figure 3F).

## Discussion

We developed a simple-to-implement pipeline for long-term imaging and tracking of double emulsion (DE) droplets. Our method addresses two common challenges in droplet-based technologies: (i) maintaining droplet position within a field of view during extended imaging assays, and (ii) the need for custom tracking software.

We prevented excessive droplet shifting by using glass-bottom dishes holding an abundance of buffer sufficient to minimize impacts of evaporation. We also took advantage of an inherent property of DE-droplets, an aqueous outer phase, to implement an optional agarose-based trapping step, further stabilizing droplet positions against accidental jostling. We conducted all image processing and data analysis with free-to-use and open source software (39).

To establish our method, we demonstrated that either approach can reduce droplet shifting within a given field of view (FOV) for up to 12 hours. Next, we showed that for droplets containing bacteria we could control the density of cells within droplets by adjusting the optical density (OD) of input cultures. We observed apparent Poissonian distributions of cells per droplet, suggesting our simple single-plane imaging method enables quantitation of bacteria approximating single-cell resolution. Lastly, we leveraged our method to monitor the growth of bacteria within many droplets in parallel, revealing heterogeneity in growth rates.

Previous droplet-monitoring efforts have either neglected individual droplet identity, or have required complex secondary trap-devices, which increase complexity and cost while reducing throughput (25,40–43); space occupied by structural traps reduces droplet density in the FOV. Our method is compatible with fully packed FOVs, maximizing throughput while eliminating the need to build specialized secondary devices. Our simple imaging set-up includes the use of widely accessible ImageJ software. While previous methods have successfully tracked single-emulsion droplets using ImageJ (2), our pipeline is the first to implement standard tracking software for DE-droplets in the absence of a dedicated trapping device. The benefits of our image analysis and tracking method also include reduced data analysis time (minutes) compared to some previously reported methods (days). (31)

Our pipeline has inherent limitations. First, our method is not designed to track individual cells over time but instead tracks total cell growth within many individual droplets in parallel. Moreover, the bulk arrangement of droplets within a sample hinders multiplexed treatment testing. Although built-in dividers allowed us to test four agarose concentrations in parallel within a single dish (Supp. Fig. 2), multiplexed condition testing remains low throughput. Furthermore, the agarose-free implementation of our method is vulnerable to mass shifting induced by accidental bumping or shaking of the microscope stage or dish during imaging. We were also limited in our capacity to interpret the observed decrease in both pixel intensity and growth rate during late timepoints; we hypothesize that photo-bleaching or cell-death during stationary phase may be contributing factors.

Our method is theoretically limited by the number of droplets that can be packed within a given FOV (∼1,500 40mm-wide droplets at 20X magnification) and the number of FOV’s imaged between timepoints. Using modified microscope stages capable of holding additional dishes, or the use of glass-bottom dishes with more dividers could help increase the number of conditions tested in parallel. Performing Z-stack imaging at high magnification may provide an opportunity to detect and track individual cells over time.

Studying individuals comprising a population is particularly exciting for phenotyping large variant libraries. Ensuring encapsulation of single entities within droplets through Poisson statistics requires working with input densities yielding mostly empty droplets. Reducing the percentage of droplets carrying two or more molecules to roughly 1% requires ∼86% of all droplets to be empty. As a result, only ∼14% of droplets are available for single-entity observations. This “Poissonian poverty” suggests that active methods for loading individuals into droplets could be useful.

Our work addresses a need for further scaling of droplet monitoring methods, which will enable a deeper understanding of the individual biological and molecular behaviors comprising population-level observations. Our results showcase the potential of simplified methods as drivers in the improvement and adoption of droplet microfluidic technologies for high-throughput phenotyping. Only by seeing the behavior of individuals comprising populations can we hope to find individual specimens satisfying specific needs and realize a more complete understanding of populations as a whole.

## Methods

Extended methods are available in the Supplemental Online Material associated with this manuscript.

### Double-Emulsion Droplet Generation

We generated double emulsion droplets in PDMS microfluidic devices using previously reported architectures(24,28). Our device featured four inlets: one for the outer aqueous phase (inlet #4), one for the oil phase (inlet #1), and two for the inner aqueous phase (inlets #2 and 3). For the initial agarose trapping optimization, we generated “empty” droplets carrying PBS pH 7.4 (Gibco) + Tween-20 (Fisher) with an oil shell of HFE7500 + 5% PicoSurf (Sphere Fluidics). Using blunt-tip needles (Millipore Sigma), we connected 1 mL plastic syringes (BD) carrying the inner solutions to LDPE medical tubing (Scientific Commodities Inc.) that were directly connected to the PDMS device. Syringe pumps (Harvard Instruments) controlled the flow of reagents into the device, with standard flow rates of 2 × 25 μL/h for two identical inner solutions of PBS (Gibco) with 1% (w/v) Tween-20 (Fisher), 125 μL/h for the fluorocarbon oil phase with 5% PicoSurf in HFE7500 (Sphere Fluidics), and 3500–3700 μL/h for the outer sheath solution of PBS with 1% (w/v) Tween-20 and 2% (w/v) Pluronic F68 (Millipore Sigma). To encapsulate bacteria in droplets, we introduced a suspension of bacteria in LB medium through the first inlet, and sterile LB medium through the second inlet at equal flow rates. To slow bacterial growth until imaging, we stored droplets on ice or at 4°C for up to 1 hour.

### Agarose Preparation and Droplet trapping

After droplet generation, we fully dissolved low-temp melt agarose powder (Invitrogen) at the indicated concentrations in outer phase solution (PBS or LB with 1% (w/v) Tween-20 and 2% (w/v) Pluronic F68 (Millipore Sigma)). We fully dissolved the agarose solutions and kept at 45°C until the trapping began. First, we used a pipette to deposit a droplet sample at the bottom of an agarose-filled, glass-bottom dish (Cellvis). We then gently swirled the dish to ensure the droplets settled on top of the glass surface as a highly confluent monolayer before agarose solidification. We allowed the dish to sit at room temperature for 5 minutes before introducing it into a pre-warmed, temperature controlled microscope at 37°C for long-term imaging.

### Strains and Growth Conditions

We generated the input bacterial stocks used throughout our experiments by inoculating E.coli carrying an mCherry + Kanamacyn resistance cassette from a glycerol stock into LB+Kan. The inoculate grew at 37°C, shaking at 250 RPM overnight to saturation. We placed the cultures on ice for 10 minutes before measuring their OD. Next, we placed 1 mL aliquots into 1.5 mL eppendorf tubes, and spinned down at 1000 x g in a chilled centrifuge (4°C) for 5 minutes. We discarded the supernatant, then we washed and re-suspended the pellet with a volume of fresh LB that would yield the desired dilution or concentration factor. Careful up-and-down pipetting broke apart cell clumps from the pellet and avoided clogging the microfluidic device.

### Microscopy & Image Processing

We captured brightfield and fluorescence images on an inverted light microscope (Nikon) with an Oko temperature control chamber set to 37°C. We obtained image stacks by imaging several fields of view within a given sample in five minute intervals. We performed image processing using FIJI software. We used the background subtraction function for flat-field correction, using a 25 pixel rolling-ball. We performed droplet detection, tracking and fluorescence quantification, using the TrackMate plugin, maximizing the number of detected droplets while minimizing false positives (Supp. Fig. 4). We used custom python scripts for all downstream data analysis and plotting.

## Author contributions

SSC, SMT, and DE conceived the project. SSC and SMT prepared all experimental samples. SSC performed all microscopy, analyzed data, prepared figures, and wrote the manuscript with input from all authors. SMT, PMF, and DE provided funding and supervision. All authors provided manuscript edits and have given approval to the final version of the manuscript.

## Acknowledgments

We thank Dr. Youngbin Lim for help establishing the microscopy pipeline and Matt DeJong for assistance with FIJI-based analysis. We are especially grateful to Prof. Paul Bollyky and Dr. Jessica Sacher for valuable discussions and phage strains, and to Prof. Brian Hie, Samuel King, Kelly Ann Lynch, and the Jacobs-Wagner lab for sharing *E. coli* strains and fluorescent reporters. This work was supported by a Garden Grant from the Homeworld Collective awarded to SMT and an NIH DP1CA290563 awarded to PMF. SSC is supported by the NSF GRFP. SMT is supported by the Arnold and Mabel Beckman Foundation through the Arnold O. Beckman Postdoctoral Fellowship in Chemical Sciences. PMF is a Chan Zuckerberg Biohub Investigator. Figure 1D was created with **BioRender**.

## Abbreviations

CPD: cells per droplet
DE: double emulsion
FOV: field of view
IQR: interquartile range
LDPE: low density polyethylene
OD: optical density
PBS: phosphate buffered saline
PDF: probability density function
PDMS: polydimethylsiloxane
SE: single emulsion

## Supporting Information for

Affordable Immobilization of Double Emulsion for High-Throughput Time-Lapse Imaging

## Supplementary Files

Supplementary files are separate files available at https://osf.io/4axv5/. All data are paired with any custom python scripts used for analysis.

**Data Supplement 1** is time-lapse microscopy data (.tif) used for analysis in Figure 1.

**Data Supplement 2** is CSV files of droplet tracks (.csv) used for analysis in Figure 1.

**Data Supplement 3** is static microscopy data (.tif) used for analysis in Figure 2.

**Data Supplement 4** is time-lapse microscopy data (.tif) used for analysis in Figure 3.

**Data Supplement 5** is a CSV file of droplet tracks (.csv) used for analysis in Figure 3.

## Supplementary Tables

Supplementary Table S1 is included below

Table S1 contains source data for observed cell counts in low-density droplets from figure 2C.

Supplementary Table S2 is included below

Table S2 contains data regarding growth rates and IQRsused to generate supplementary figure 4

**Table S1.** Observed Cell Counts for Low-density Droplets.

| **Cell Number** | **0** | **1** | **2** | **3** | **4+** |
| --- | --- | --- | --- | --- | --- |
| **Count** | 28 | 40 | 29 | 10 | 4 |

**Table S2.** Mean Growth Rate Across All Droplets and IQR for Each Timepoint.

| **Time (Minutes)** | **Mean Growth Rate (Doublings/Hour)** | **IQR** |
| --- | --- | --- |
| 15.0 | 0.04 | 0.25 |
| 25.0 | 0.07 | 0.27 |
| 35.0 | 0.12 | 0.24 |
| 45.0 | 0.21 | 0.26 |
| 55.0 | 0.35 | 0.24 |
| 65.0 | 0.46 | 0.24 |
| 75.0 | 0.56 | 0.25 |
| 85.0 | 0.61 | 0.25 |
| 95.0 | 0.65 | 0.27 |
| 105.0 | 0.71 | 0.21 |
| 115.0 | 0.72 | 0.25 |
| 125.0 | 0.78 | 0.24 |
| 135.0 | 0.8 | 0.21 |
| 145.0 | 0.81 | 0.22 |
| 155.0 | 0.82 | 0.23 |
| 165.0 | 0.8 | 0.22 |
| 175.0 | 0.76 | 0.24 |
| 185.0 | 0.7 | 0.22 |
| 195.0 | 0.61 | 0.22 |
| 205.0 | 0.52 | 0.21 |
| 215.0 | 0.43 | 0.25 |
| 225.0 | 0.34 | 0.22 |
| 235.0 | 0.27 | 0.16 |
| 245.0 | 0.19 | 0.18 |
| 255.0 | 0.14 | 0.17 |
| 265.0 | 0.09 | 0.16 |
| 275.0 | 0.04 | 0.15 |
| 285.0 | 0.0 | 0.14 |
| 295.0 | 0.0 | 0.13 |
| 305.0 | -0.04 | 0.14 |
| 315.0 | -0.07 | 0.11 |
| 325.0 | -0.07 | 0.11 |
| 335.0 | -0.08 | 0.11 |
| 345.0 | -0.1 | 0.11 |
| 355.0 | -0.11 | 0.1 |
| 365.0 | -0.11 | 0.1 |
| 375.0 | -0.1 | 0.1 |

## Supplementary Figures

**Supplementary Figure 1:**
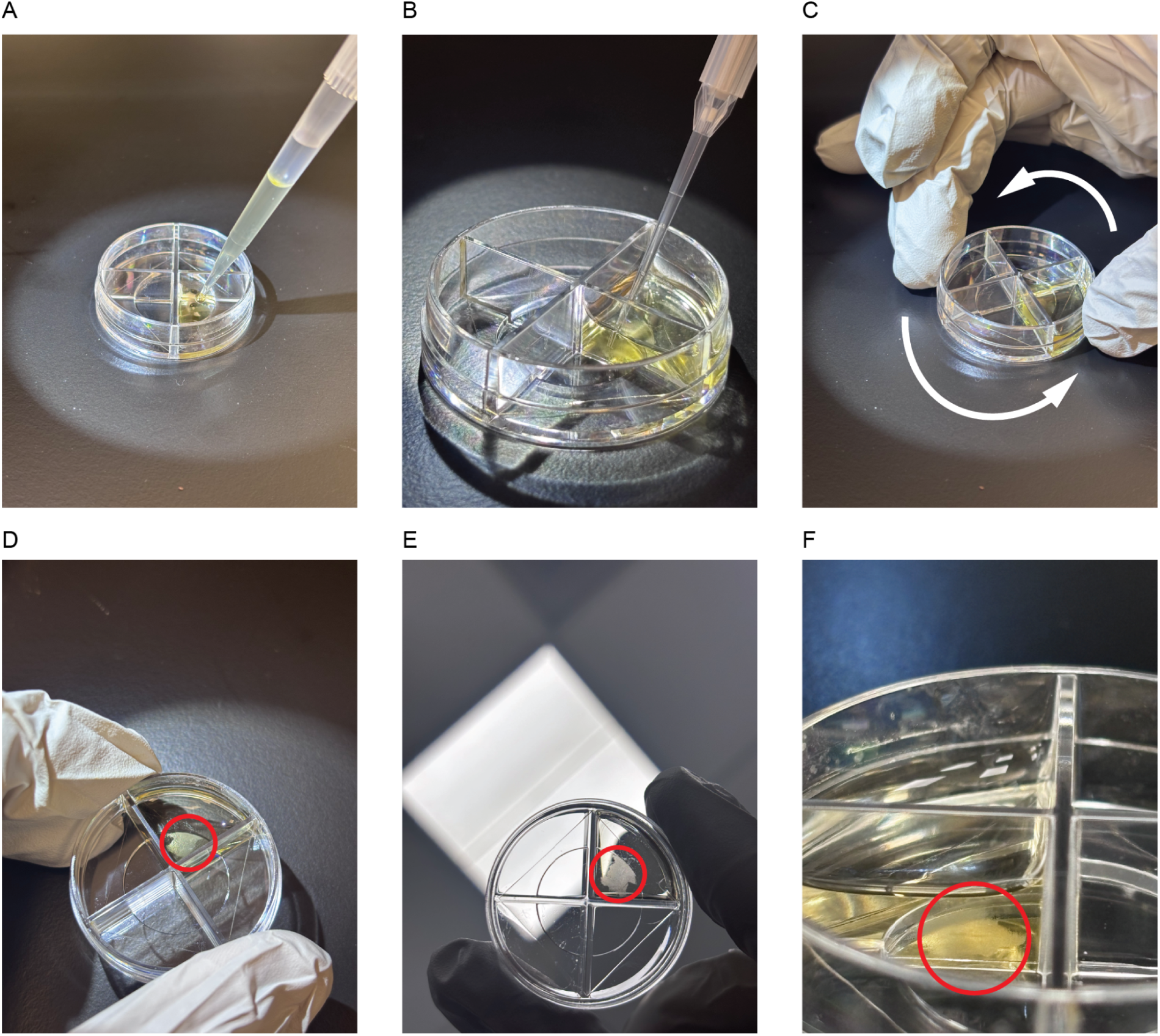
Droplet trapping protocol. (a) 400 µL of melted agarose at 45°C are pipetted into a glass-bottom dish. (b) Droplets are injected into the bottom of the dish while agarose is still liquid (c) Dish is swirled to help droplets fall to the glass surface and rest as a confluent monolayer before agarose solidification. (d),(e),(f) shows successfully trapped droplets from different angles. Red circles mark the droplet monolayer.

**Supplementary Figure 2:**
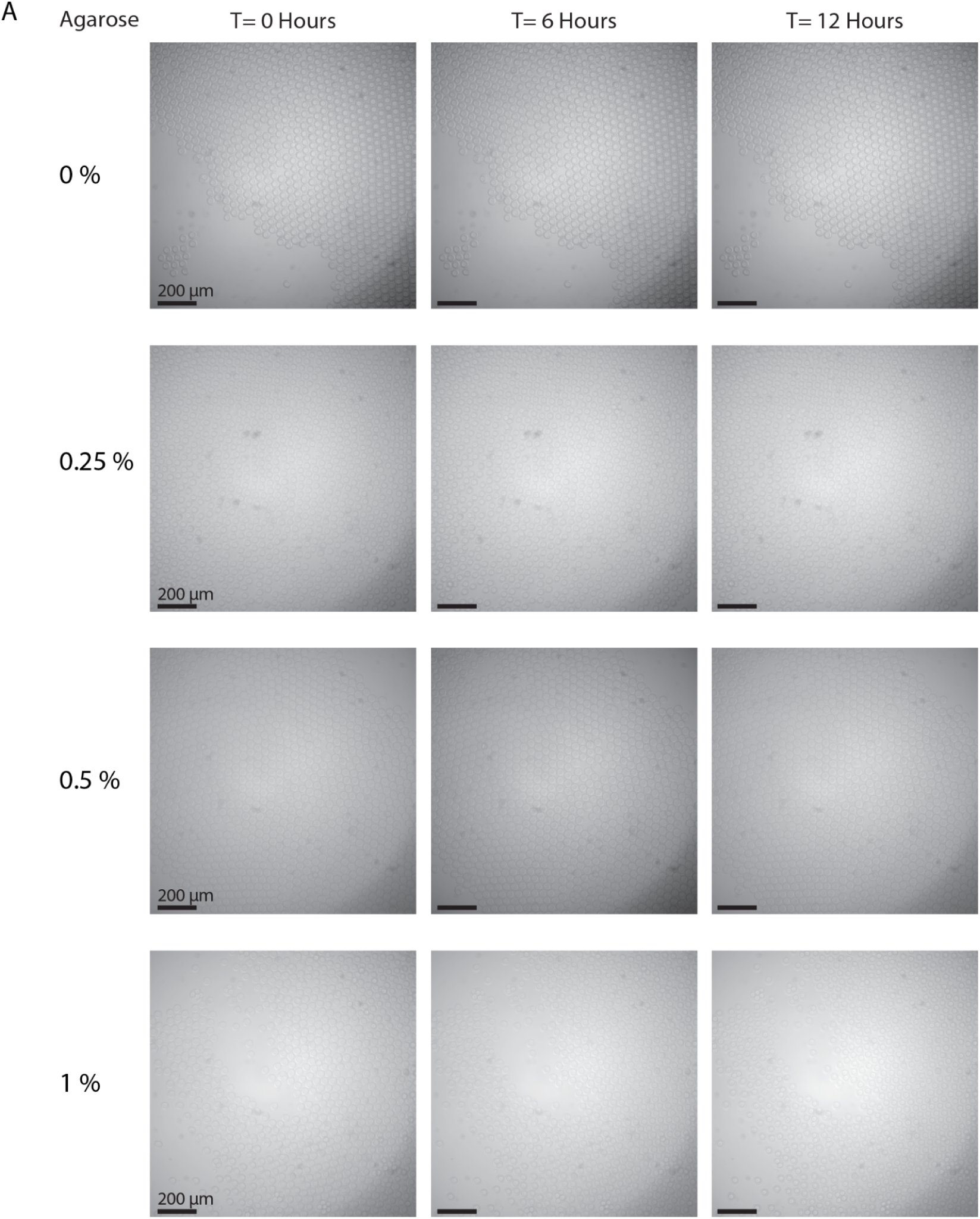
Microscopy shows droplets are compatible with agarose embedding for up to 12 hours. (a) Droplets fixed in PBS-containing agarose at concentrations ranging from 0 to 1% W/V. All conditions were tested in a glass-bottom dish using 400uL of outer buffer. Each row represents a different agarose concentration. Each column represents a unique timepoint.

**Supplementary Figure 3:**
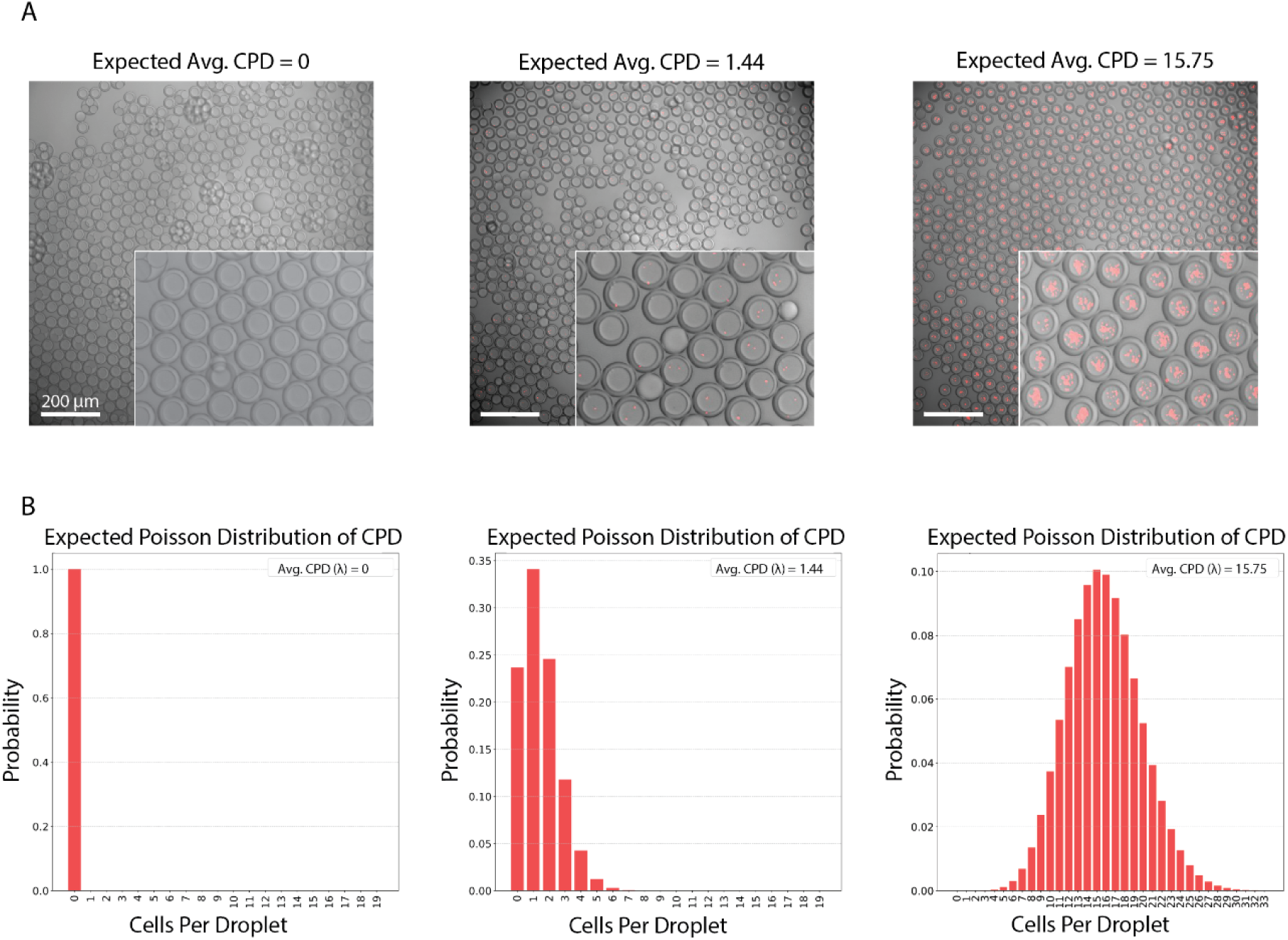
Microscopy of empty and high-density droplets tracks expected Poisson distributions. (a) Merged brightfield and fluorescence images of droplets at different cell-density regimes (left to right: OD_600_ of 0, 0.86, and 9.4). (b) Expected Poisson distribution of cell occupancy within droplets for each cell density regime.

**Supplementary Figure 4:**
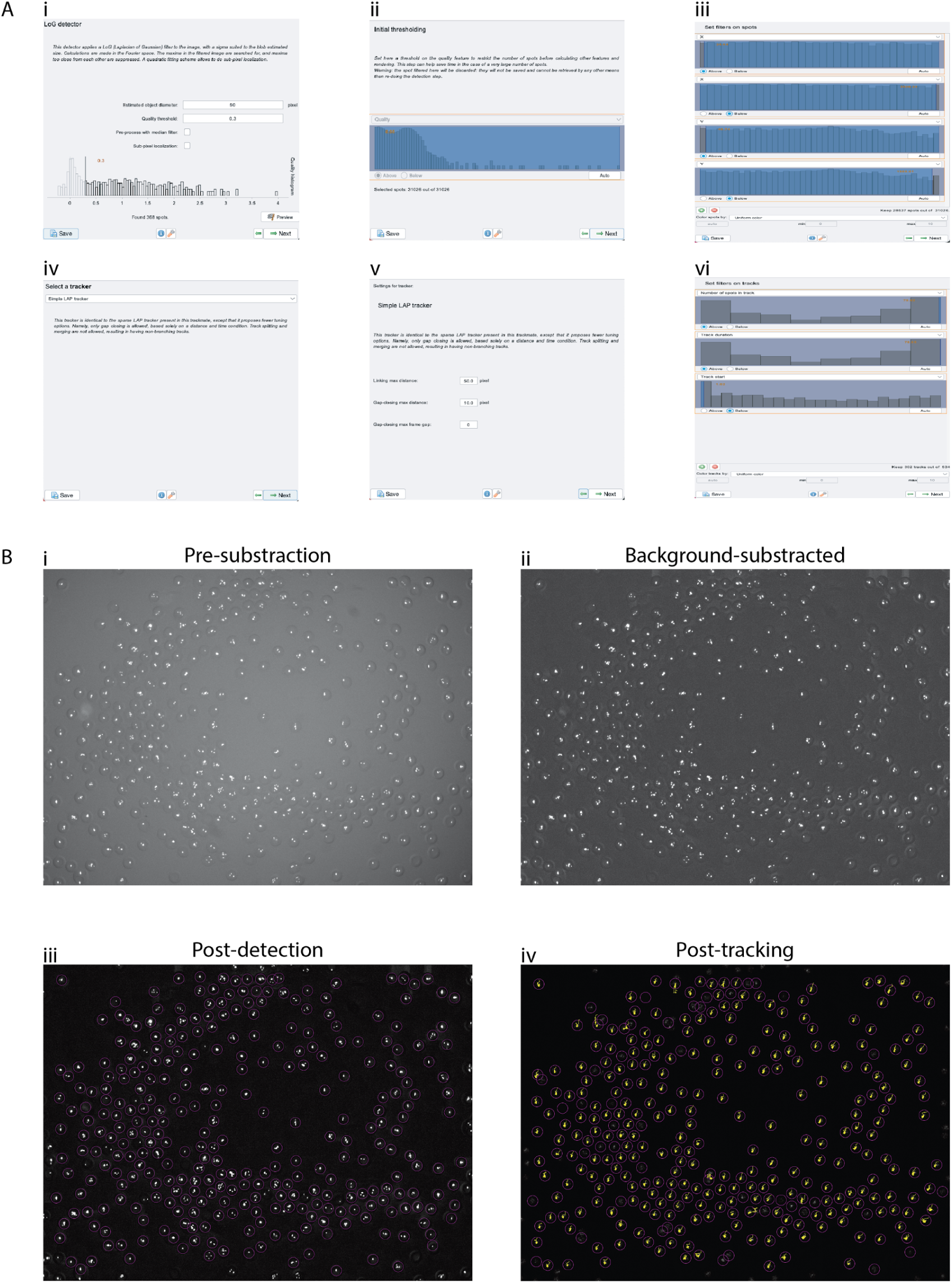
FIJI data analysis pipeline. (a) Steps for droplet detection and tracking in FIJI within the graphic user interface (GUI). (b) Images depicting progression of hyperstack before and during analysis using TrackMate’s GUI. Images i and ii are obtained before executing TrackMate with FIJI tools.

## Operating microfluidic devices

### Preparing stock reagents

For the inner solutions, we used either PBS (Gibco, cat. # 10010031) at pH 7.4 or cell cultures in LB (Difco, BD, cat. # 244620) with Kanamycin (50 µg/mL, Gibco, cat #15160054) at the desired OD. In either case, we added Tween 20 (Fisher BioReagents, cat. # BP337) at 1% by measuring 1 gram of Tween 20 in a graduated cylinder and bringing the volume up to 100 mL with each solvent. We adjusted the cell cultures to the desired OD by centrifugation at 10,000 RCF for 2 minutes, followed by pellet washing and re-suspension in fresh LB media. To prevent cellular clumping, we added 1% BSA to the fresh LB used in washing/re-suspension steps. For the outer solution, we mixed 1% Tween 20 + 2% Pluronic F68/Kolliphor P 188 (Millipore Sigma, cat. # K4894) by adding 1 g of Tween 20 and 2 g of Pluronic F68 into a graduated cylinder and bringing the volume up to 100 mL with PBS pH 7.4 or fresh LB + Kanamycin. To each graduated cylinder, we added a stir bar and the opening was sealed with Parafilm (Bemis, cat. # PM-996). We dissolved the surfactants by overturning and stirring, and later sterile-filtered the solutions into fresh PETG storage bottles (Nalgene, cat. # 342020-0030).

### PDMS device generation

We used PDMS devices found in previous reports [16, 23]. CAD files for photolithography exposure masks are available from the Open Science Framework (OSF) repository for that reference: https://osf.io/h4sr9/. Briefly, we generated molds for device features on silicon wafers using photolithography. We then casted PDMS (MG Chemicals, cat. # RTV615) onto a feature mold and a blank silicon water; after a soft bake (80°C, 12-15 min), we released the PDMS from the mold. We punched holes for inlet and outlet ports with a 1 mm biopsy punch (Robbins Instruments, cat. #RBP-10), cleaned the PDMS pieces with Scotch tape, and joined the featured and blank PDMS slabs before baking them at 80°C for 48-56 hrs. AutoCAD files and photolithography protocols for masks used in this study are available in the following OSF repository: https://osf.io/4axv5/.

### Generating double emulsion droplets

We generated double emulsions as previously described using a set-up that included the PDMS devices, 4 syringe pumps (Harvard Apparatus, cat. # 70-4511), and a zoom power optical microscope. We prepared aqueous solutions as described above and filtered with a syringe-driven 0.22 µm filter. We used 5% PicoSurf in HFE7500 (Sphere Fluidics) as the droplet oil for the fluorinated oil shell.

For DE generation, we loaded input solutions into disposable Luer lock syringes: inner solution in two 1 mL syringes, PicoSurf oil in a 3 mL or 5 mL syringe, and outer solution in a 10 mL syringe (Becton Dickinson, cat. # 309628, 309657, 309646, 302995). We removed bubbles from each solution by tapping and ejection, then capped each syringe with a 27-gauge stainless steel dispensing needle (Millipore Sigma, cat. # 917532-1EA). We inserted the needles into the open end of polyethylene medical tubing (Scientific Commodities Inc., cat. # BB31695-PE/2). After installing the syringes into the pumps, we cut the tubing with sufficient length to reach the PDMS device on the stage. We centered the device in the microscope field of view, and inserted the free ends of the tubing directly into the ports of the PDMS device after trimming the tubing to reduce slack. We removed the tubing, and plasma treated the right-hand side of the device to render it hydrophilic. We covered the ports for inner and oil phases with Scotch tape, and the PDMS device array was placed in a plasma cleaner (Harrick Plasma, cat. # PDC-001) with a dry scroll pump (Agilent cat. # IDP3B01), a Type 0536 TC vacuum gauge (Agilent cat. # L6141303), and a vacuum gauge monitor (Harrick Plasma, cat. # PDCVCG). We vacated the chamber, and opened the three-way valve to ambient air with the regulator valve that maintained pressure at approximately 400 mbar. We performed plasma treatment under a “high” setting (30 watts) for 10-12 minutes. After deactivating plasma coils, we repressurized the chamber to ambient pressure and removed the PDMS device array. During plasma treatment, we ejected sufficient fluid solutions to fill the tubing with solution and eliminate air bubbles. After plasma treatment, we operated 30 µm devices as follows. We connected an outlet line to the device and the free end was inserted into a fresh 2 mL tube (port #4 on figure 1A) After centering the device on the microscope, and connecting the outer solution line to the device (port #5), we flowed the outer solution through the port for 30 sec at a flow rate of 3000 µL/hr. We set the oil input (port #3) to a flow rate of 400 µL/hr and then connected the tubing to the port. Once the oil was visible in the second first flow focuser we cut back the flow rate to 200 µL/hr and progressively adjusted to 125 µL/hr. Finally, we set the two inner inputs to flow rates of 100 µL/hr each and connected the lines to the device (ports #2 and 3). When the inner solution was visible in the first flow focuser, we decreased each flow rate to 25 µl/hr. We gradually increased the outer flow rate to 3500 µL/hr. We periodically adjusted flow to maintain stable DE generation or to alter the DE geometry.

### Imaging double emulsion droplets

#### Microscopy

We imaged DEs on an inverted light microscope (Nikon, Eclipse Ti). We observed robust integrity of generated droplets when kept on ice for up to 72 hours pre-imaging. We resuspended samples consisting of double emulsions in solution by overturning the sample tube, and then pipetted 10 µL of undiluted solution onto a Countess chamber slide (ThermoFisher Scientific, cat. # C10228) without agarose or into a glass-bottom dish (Cellvis, cat. # D35C4-20-0-N) with and without agarose. In the case of the glass-bottom dish, we pipetted the 10 uL of droplet sample into 400uL of outer solution to minimize evaporation-induced droplet shifting. We controlled the light microscope with a distribution of the MicroManager software package. We imaged with a combination of brightfield (Semrock, BRFLD-A-NTE-ZERO) and red fluorescence (Semrock,LED-mCherry-A-NTE-ZERO). Exposure times were 10 ms for bright field and 500 ms for red fluorescence (Mcherry1). A SOLA SE light engine provided illumination through a 3mm liquid light guide (Lumencor, cat # SOLA SE 5-LCR-SA). We used a 10x objective lens for all images (Nikon, cat# MRD00100). We captured images on an sCMOS camera (Andor, Zyla 4.2 Plus, VSC-06278). We recorded all images at 1×1 binning. We analyzed DE images with FIJI and custom Python scripts. Scripts and data are available at https://osf.io/4axv5/.

#### FIJI analysis pipeline

Briefly, we loaded image stacks (.tif) into Fiji and created the relevant channel sub-stack (brightfield-only for agarose optimization, fluorescence-only for growth tracking). We first processed the sub-stacks through a “background subtraction” step akin to flatfield-correction, using a 25 pixel rolling-ball radius. We then analyzed the normalized sub-stack images via Fiji’s Trackmate Plugin. We selected the LoG (Laplacian of Gaussian) detector for the initial detection step. We selected an estimated DE diameter of 50 pixels. For the spot detection threshold, we chose a quality score of 0.3 or above to filter out false positives. To further filter the spots, we adjusted the X and Y position limits to avoid recording data from droplets at the edge of our field-of-view.

For tracking, we used the “Simple LAP tracker” function, setting the linking max distance to 50 pixels. We set the “gap-closing max distance” to 10 pixels and the “gap-closing max frame gap” to 0. We next set filters on tracks based on the number of spots in track, track duration, and track start frame. We only kept tracks that followed a spot present from frame 0 to frame 80, without gaps in detection. We extracted tables containing information about the track lengths and the spot intensities as .CSV files and used them for downstream analysis in Python using “Mean Intensity” as the main metric for fluorescence.

## Sources

1. Arter WE, Qi R, Erkamp NA, Krainer G, Didi K, Welsh TJ, et al. Biomolecular condensate phase diagrams with a combinatorial microdroplet platform. Nat Commun. 2022 Dec 21;13(1):7845. doi:10.1038/s41467-022-35265-7

2. Pratt SL, Zath GK, Akiyama T, Williamson KS, Franklin MJ, Chang CB. DropSOAC: Stabilizing Microfluidic Drops for Time-Lapse Quantification of Single-Cell Bacterial Physiology. Front Microbiol. 2019 Sep 24;10. doi:10.3389/fmicb.2019.02112

3. Qu F, Zhao S, Cheng G, Rahman H, Xiao Q, Chan RWY, et al. Double emulsion-pretreated microwell culture for the in vitro production of multicellular spheroids and their in situ analysis. Microsyst Nanoeng. 2021 May 24;7(1):38. doi:10.1038/s41378-021-00267-w

4. Yu JQ, Huang W, Chin LK, Lei L, Lin ZP, Ser W, et al. Droplet optofluidic imaging for λ-bacteriophage detection via co-culture with host cell Escherichia coli. Lab Chip. 2014 Aug 11;14(18):3519–24. doi:10.1039/C4LC00042K

5. Jeong HH, Hyung Jin S, Jin Lee B, Kim T, Lee CS. Microfluidic static droplet array for analyzing microbial communication on a population gradient [Internet]. 2015 Jan 22. doi:10.1039/C4LC01097C

6. Nauwynck W, Faust K, Boon N. Droplet microfluidics for single-cell studies: a frontier in ecological understanding of microbiomes. FEMS Microbiol Rev. 2025 Jan 1;49:fuaf032. doi:10.1093/femsre/fuaf032

7. Lim SW, Abate AR. Ultrahigh-throughput sorting of microfluidic drops with flow cytometry. Lab Chip. 2013 Oct 30;13(23):4563–72. doi:10.1039/C3LC50736J

8. Brower KK, Carswell-Crumpton C, Klemm S, Cruz B, Kim G, Calhoun SGK, et al. Double emulsion flow cytometry with high-throughput single droplet isolation and nucleic acid recovery. Lab Chip. 2020 Jun 16;20(12):2062–74. doi:10.1039/D0LC00261E

9. Scheele RA, Weber Y, Nintzel FEH, Herger M, Kaminski TS, Hollfelder F. Ultrahigh Throughput Evolution of Tryptophan Synthase in Droplets via an Aptamer Sensor. ACS Catal. 2024 Apr 19;14(8):6259–71. doi:10.1021/acscatal.4c00230

10. Bernath K, Hai M, Mastrobattista E, Griffiths AD, Magdassi S, Tawfik DS. In vitro compartmentalization by double emulsions: sorting and gene enrichment by fluorescence activated cell sorting. Anal Biochem. 2004 Feb 1;325(1):151–7. doi:10.1016/j.ab.2003.10.005

11. Sukovich DJ, Lance ST, Abate AR. Sequence specific sorting of DNA molecules with FACS using 3dPCR. Sci Rep. 2017 Jan 4;7(1):39385. doi:10.1038/srep39385

12. Gupta RD, Goldsmith M, Ashani Y, Simo Y, Mullokandov G, Bar H, et al. Directed evolution of hydrolases for prevention of G-type nerve agent intoxication. Nat Chem Biol. 2011 Feb;7(2):120–5. doi:10.1038/nchembio.510

13. Lance ST, Sukovich DJ, Stedman KM, Abate AR. Peering below the diffraction limit: robust and specific sorting of viruses with flow cytometry. Virol J. 2016 Dec 1;13(1):201. doi:10.1186/s12985-016-0655-7

14. Ma F, Xie Y, Huang C, Feng Y, Yang G. An Improved Single Cell Ultrahigh Throughput Screening Method Based on In Vitro Compartmentalization. PLOS ONE. 2014 Feb 24;9(2):e89785. doi:10.1371/journal.pone.0089785

15. Holstein JM, Gylstorff C, Hollfelder F. Cell-free Directed Evolution of a Protease in Microdroplets at Ultrahigh Throughput. ACS Synth Biol. 2021 Feb 19;10(2):252–7. doi:10.1021/acssynbio.0c00538

16. Rosenthal RG, Diana Zhang X, Đurđić KI, Collins JJ, Weitz DA. Controlled Continuous Evolution of Enzymatic Activity Screened at Ultrahigh Throughput Using Drop-Based Microfluidics. Angew Chem Int Ed. 2023;62(24):e202303112. doi:10.1002/anie.202303112

17. Stucki A, Vallapurackal J, Ward TR, Dittrich PS. Droplet Microfluidics and Directed Evolution of Enzymes: An Intertwined Journey. Angew Chem Int Ed. 2021;60(46):24368–87. doi:10.1002/anie.202016154

18. Aubermann F, Seneca S, Hofman T, Garcés-Lázaro I, Ajmail K, Daubner K, et al. Modular Droplet-Based Microfluidic Platform for Functional Phenotypic Screening of Natural Killer Cells. Small Methods. 2025;9(8):2500236. doi:10.1002/smtd.202500236

19. Jain A, Stavrakis S, deMello A. Droplet-based microfluidics and enzyme evolution. Curr Opin Biotechnol. 2024 Jun 1;87:103097. doi:10.1016/j.copbio.2024.103097

20. Manteca A, Gadea A, Van Assche D, Cossard P, Gillard-Bocquet M, Beneyton T, et al. Directed Evolution in Drops: Molecular Aspects and Applications. ACS Synth Biol. 2021 Nov 19;10(11):2772–83. doi:10.1021/acssynbio.1c00313

21. Han HS, Cantalupo PG, Rotem A, Cockrell SK, Carbonnaux M, Pipas JM, et al. Whole-Genome Sequencing of a Single Viral Species from a Highly Heterogeneous Sample. Angew Chem Int Ed. 2015;54(47):13985–8. doi:10.1002/anie.201507047

22. Clausell-Tormos J, Lieber D, Baret JC, El-Harrak A, Miller OJ, Frenz L, et al. Droplet-Based Microfluidic Platforms for the Encapsulation and Screening of Mammalian Cells and Multicellular Organisms. Chem Biol. 2008 May 19;15(5):427–37. doi:10.1016/j.chembiol.2008.04.004

23. Liu H, Zhang Y, Kroukamp H, Peng K, Cain AK, Paulsen IT, et al. Screening and selection of cellulase-secreting yeast single cells using integrated double emulsion droplet and flow cytometry techniques. Sens Actuators B Chem. 2024 Oct 1;416:136038. doi:10.1016/j.snb.2024.136038

24. McCully AL, Loop Yao M, Brower KK, Fordyce PM, Spormann AM. Double emulsions as a high-throughput enrichment and isolation platform for slower-growing microbes. ISME Commun. 2023 Dec 1;3(1):47. doi:10.1038/s43705-023-00241-9

25. Barizien A, Suryateja Jammalamadaka MS, Amselem G, Baroud CN. Growing from a few cells: combined effects of initial stochasticity and cell-to-cell variability. J R Soc Interface. 2019 Apr 24;16(153):20180935. doi:10.1098/rsif.2018.0935

26. Huang S, Srimani JK, Lee AJ, Zhang Y, Lopatkin AJ, Leong KW, et al. Dynamic control and quantification of bacterial population dynamics in droplets. Biomaterials. 2015 Aug 1;61:239–45. doi:10.1016/j.biomaterials.2015.05.038

27. Yang Z, Thompson S, Zhang Y, Rutten I, Van Duyse J, Van Isterdael G, et al. Continuous FACS Sorting of Double Emulsion Picoreactors with a 3D-Printed Vertical Mixer. Anal Chem. 2025 Jul 15;97(27):14406–14. doi:10.1021/acs.analchem.5c01536

28. Thompson S, Zhang Y, Yang Z, Nichols L, Fordyce PM. FACS-Sortable Triple Emulsion Picoreactors for Screening Reactions in Biphasic Environments. Adv Mater Interfaces. 2025;12(3):2400403. doi:10.1002/admi.202400403

29. Fergola A, Ballesio A, Frascella F, Napione L, Cocuzza M, Marasso SL. Droplet Generation and Manipulation in Microfluidics: A Comprehensive Overview of Passive and Active Strategies. Biosensors. 2025 Jun;15(6):345. doi:10.3390/bios15060345

30. Sabhachandani P, Sarkar S, Zucchi PC, Whitfield BA, Kirby JE, Hirsch EB, et al. Integrated microfluidic platform for rapid antimicrobial susceptibility testing and bacterial growth analysis using bead-based biosensor via fluorescence imaging. Microchim Acta. 2017 Dec 1;184(12):4619–28. doi:10.1007/s00604-017-2492-9

31. Taylor D, Verdon N, Lomax P, Allen RJ, Titmuss S. Tracking the stochastic growth of bacterial populations in microfluidic droplets. Phys Biol. 2022 Feb;19(2):026003. doi:10.1088/1478-3975/ac4c9b

32. Amselem G, Guermonprez C, Drogue B, Michelin S, N. Baroud C. Universal microfluidic platform for bioassays in anchored droplets [Internet]. 2016 Oct 18. doi:10.1039/C6LC00968A

33. Sart S, Tomasi RFX, Amselem G, Baroud CN. Multiscale cytometry and regulation of 3D cell cultures on a chip. Nat Commun. 2017 Sep 7;8(1):469. doi:10.1038/s41467-017-00475-x

34. Sabhachandani P, Motwani V, Cohen N, Sarkar S, Torchilin V, Konry T. Generation and functional assessment of 3D multicellular spheroids in droplet based microfluidics platform. Lab Chip. 2016 Jan 26;16(3):497–505. doi:10.1039/C5LC01139F

35. McMillan KS, McCluskey AG, Sorensen A, Boyd M, Zagnoni M. Emulsion technologies for multicellular tumour spheroid radiation assays. Analyst. 2015 Dec 14;141(1):100–10. doi:10.1039/C5AN01382H

36. Huebner A, Bratton D, Whyte G, Yang M, J. deMello A, Abell C, et al. Static microdroplet arrays: a microfluidic device for droplet trapping, incubation and release for enzymatic and cell-based assays. Lab Chip. 2009;9(5):692–8. doi:10.1039/B813709A

37. Gerver RE, Gómez-Sjöberg R, Baxter BC, Thorn KS, Fordyce PM, Diaz-Botia CA, et al. Programmable microfluidic synthesis of spectrally encoded microspheres. Lab Chip. 2012 Oct 16;12(22):4716–23. doi:10.1039/C2LC40699C

38. Elowitz MB, Leibler S. A synthetic oscillatory network of transcriptional regulators. Nature. 2000 Jan;403(6767):335–8. doi:10.1038/35002125

39. Tinevez JY, Perry N, Schindelin J, Hoopes GM, Reynolds GD, Laplantine E, et al. TrackMate: An open and extensible platform for single-particle tracking. Methods San Diego Calif. 2017 Feb 15;115:80–90. doi:10.1016/j.ymeth.2016.09.016 PubMed PMID: 27713081.

40. Leung K, Zahn H, Leaver T, Konwar KM, Hanson NW, Pagé AP, et al. A programmable droplet-based microfluidic device applied to multiparameter analysis of single microbes and microbial communities. Proc Natl Acad Sci. 2012 May 15;109(20):7665–70. doi:10.1073/pnas.1106752109

41. Boedicker JQ, Vincent ME, Ismagilov RF. Microfluidic Confinement of Single Cells of Bacteria in Small Volumes Initiates High-Density Behavior of Quorum Sensing and Growth and Reveals Its Variability. Angew Chem Int Ed. 2009;48(32):5908–11. doi:10.1002/anie.200901550

42. Guo X, Silva KPT, Boedicker JQ. Single-cell variability of growth interactions within a two-species bacterial community. Phys Biol. 2019 Mar;16(3):036001. doi:10.1088/1478-3975/ab005f

43. Lu H, Caen O, Vrignon J, Zonta E, El Harrak Z, Nizard P, et al. High throughput single cell counting in droplet-based microfluidics. Sci Rep. 2017 May 2;7(1):1366. doi:10.1038/s41598-017-01454-4

